# Active shrinkage protects neurons following axonal transection

**DOI:** 10.1101/2022.07.06.499034

**Authors:** Mehmet Şerif Aydın, Sadık Bay, Esra Nur Yiğit, Cemil Özgül, Elif Kaval Oğuz, Elçin Yenidünya Konuk, Neşe Ayşit, Nureddin Cengiz, Ender Erdoğan, Aydın Him, Mehmet Koçak, Emrah Eroglu, Gürkan Öztürk

## Abstract

Trauma, vascular events, or neurodegenerative processes can lead to axonal injury and eventual transection (axotomy). Neurons can survive axotomy, yet the underlying mechanisms are not fully understood. Excessive water entry into injured neurons poses a particular risk due to swelling and subsequent death. Using *in vitro* and *in vivo* neurotrauma model systems based on laser transection, we demonstrated that axotomy triggers actomyosin contraction coupled with calpain activity. As a consequence, neurons shrink acutely to force water out through aquaporin channels preventing swelling and bursting. Inhibiting shrinkage increased the probability of neuronal cell death by about three-fold. These studies reveal a previously unrecognized cytoprotective response mechanism to neurotrauma and offer a fresh perspective on pathophysiological processes in the nervous system.

**One-Sentence Summary:** When the axon of a neuron is cut, its soma shrinks to pump out water to avoid deadly swelling.

## Introduction

Axons of most neurons can extend to several thousand-fold lengths of the cell’s diameter. Such a prominent structure represents a substantial cellular surface area susceptible to intrusions of various kinds, ranging from traumas to autoimmunity. Many nervous system disorders start or progress by axonal injury, which may result in the death of the affected neuron, acutely or over extended periods (*1, 2*). In the aftermath of an axon transection (axotomy), the neuron becomes susceptible to Na^+^, C^-^, and Ca^++^ influx through the cut-end until effective resealing, especially in injuries close to the neuronal body. Ion influx occurs through specific channels in the soma that open due to changes in the plasma membrane’s electrical potential or mechanical stimuli caused by the injury (*3*). Water accompanies ions into the neuron as the cytosol becomes more and more hyperosmolar (*3, 4*). Two main mechanisms are known to decrease solute load and reduce the risk of swelling after axonal injury. First, during strong depolarization triggered by an axonal injury, the Na^+^-Ca^++^ exchange pump drives out Na^+^ in exchange for Ca^++^ (*5, 6*). However, cytosolic Ca^++^-overload may damage the plasma membrane, mitochondria, and cytoskeleton, ultimately leading to cell death (*5*). To remove excessive Cl^-^ from the cytosol, cells use Cl^-^ – K^+^ cotransporters to release Cl^-^ along with K^+^ that spontaneously diffuses out due to concentration gradient. The ATP-dependent Na^+^-K^+^ pump is overactivated in response to the drop in K^+^ levels, which adds to the impending energy crisis (*4, 7, 8*).

Laser axotomy *in vitro* and living animals is an established method for examining the factors influencing neuronal survival and degenerative processes. We observed a hitherto unreported phenomenon during laser axotomy experiments: *shrinkage in neuronal soma*. This critical observation aligns with earlier findings that neurons employ local axonal contractions to promote resealing of cut axon ends (*9*). We hypothesized that actomyosin contraction (AMC) in the neuronal body might be an early response to axotomy, which counteracts the ensuing deadly swelling.

## Results & Discussion

### Axotomy causes a reduction in cell size, which is preceded by calcium influx and membrane depolarization

The *in vitro* experimental paradigm employs time-lapse imaging of adult primary sensory neurons during the transection of outgrown axons with a laser beam at physiological temperature and atmosphere. In response to laser axotomy, neurons immediately started to shrink. Cell surface area change analysis revealed that the neuronal bodies shrank about 12% within the first 15 minutes (Fig. 1A, B, and movie S1). Notably, nearly half of the shrinkage occurred during the first 5 minutes, which we selected as the reference time point for other experiments. Next, we sought to understand this observation’s *in vivo* relevance and utilized Thy1-GFP line M strain mice with sparse GFP expression in dorsal root ganglia (DRG) neurons (*10*), where individual axons can be identified, allowing precise *in vivo* laser transection. An imaging window was surgically formed to expose the second lumbar (L2) DRGs, as described before (*11*). Cutting single axons with a femtosecond infrared laser of a two-photon microscope confirmed that these neurons *in vivo* shrank comparable to isolated cells *in vitro* (Fig. 1A, B). We next axotomized hippocampal neurons in culture to determine whether axotomy-induced shrinkage also occurs in cells from the central nervous system besides primary sensory neurons. The hippocampal neurons also exhibited a significant shrinkage, albeit less than DRG neurons. We selected DRG neurons as a model system for the remainder of the study due to their excellent *in vitro* survival and the ease of intravital imaging.

**Fig. 1.**
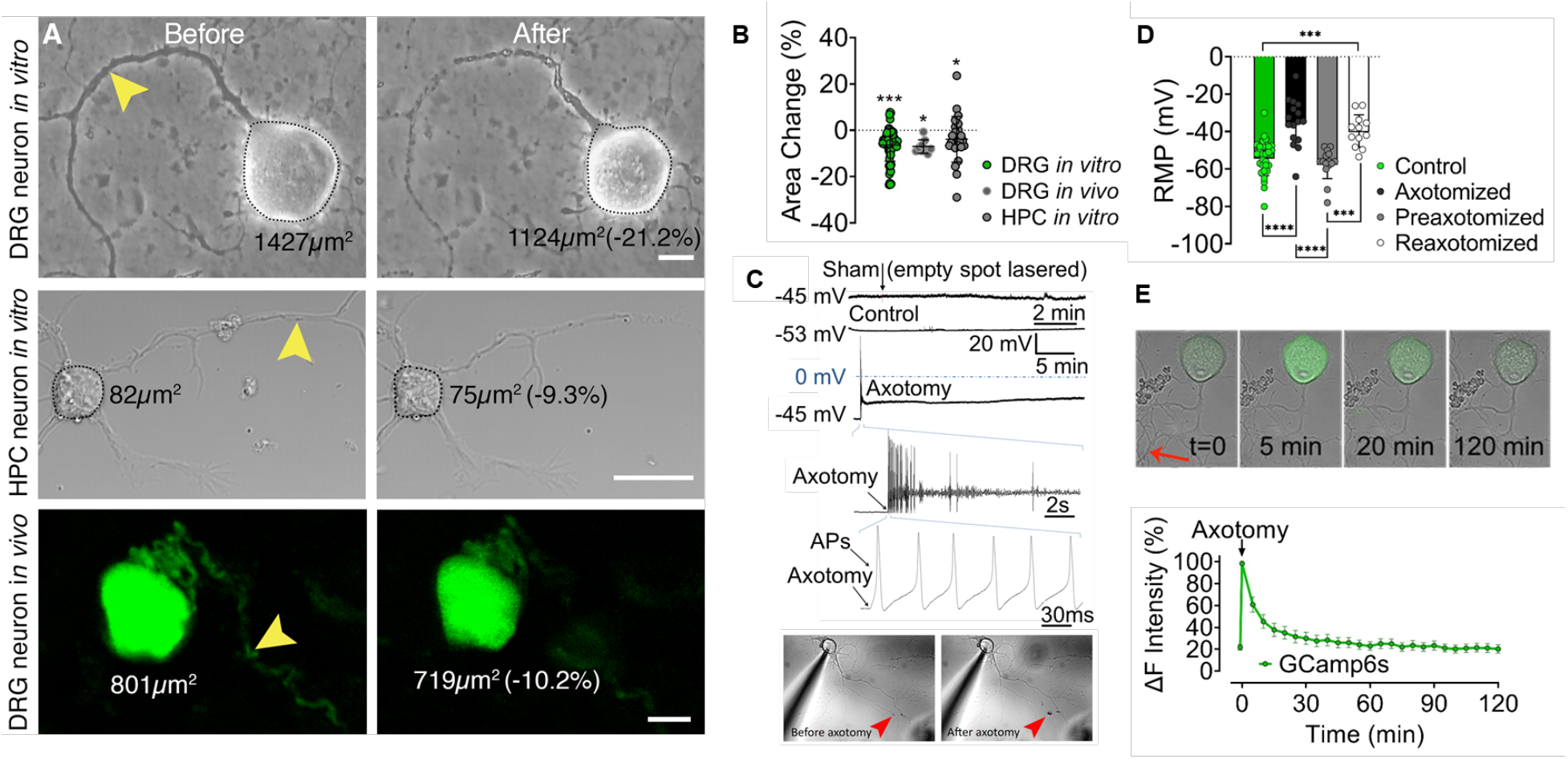
Acute consequences of axotomy. (**A**) Images show dorsal root ganglion (DRG) and hippocampal (HPC) neurons *in vitro* and a DRG neuron of a live Thy1-GFP mouse before and after laser axotomy (scale bar, 15 μm; arrowhead points to the site of axotomy). (**B**) The scatter dot plot shows changes in the surface area in 5 mins following axotomy in cultured (n=6/65) and *in vivo* DRG neurons (n=3/9) and hippocampal neurons (n=3/25). Wilcoxon Signed Ranks Test was used to compare surface areas before and after axotomy (**C**) Membrane potential recordings from a control and an axotomized neuron. Images show simultaneous patch-clamp recording and axotomy (arrowhead points to the injury site). (**D**) Bars represent levels of the resting membrane potential (RMP) of neurons before (n=4/29), immediately (n=3/16)), and 24 hrs (n=3/15) after axotomy and following the second axotomy (n=3/12); Kruskal Wallis and Mann-Whitney U tests were used to compare groups. (**E**) Real-time Ca^++^ traces upon axotomy in GCamp6-expressing neurons. Error bars: SEM; n=number of animals used/number of cells analyzed; for all tests: *p<0.05, ***p<0.0001, ****p<0.00001.

Axotomy causes membrane depolarization and action potentials in cortical neurons *in vitro* (*3*). Since many post-axotomy events can be related to this and concurrent Ca^++^ influx, we first characterized these phenomena using patch-clamp recordings. We exploited DRG neurons from genetically engineered mice expressing the genetically encoded Ca^++^ biosensor GCamp6 to capture neuronal body shrinkage and subsequent Ca^++^ influx in response to laser axotomy. Laser axotomy significantly depolarized the membrane with or without generating action potentials (AP) (Fig. 1C). After 24 hrs, the resting membrane potentials of axotomized neurons were similar to the controls, and when re-axotomized then, the membrane potential depolarized comparable to naïve neurons, but AP generation was sporadic (Fig. 1D and supplementary text). Live-cell Ca^++^ imaging showed that cytosolic Ca^++^ levels plateaued sharply after axotomy and returned to baseline within an hour (Fig. 1E and movie S2). We also observed a persistent Ca^++^ influx that led to enhanced cytosolic Ca^++^ accumulation when the proximal stump was not promptly resealed (movie S3 and S4).

### Calcium entry through voltage-gated channels triggers AMC

Ca^++^ may enter the cell through voltage-gated calcium channels (VGCCs) in addition to the cut ends of the injured axons (*5*). DRG neurons express five types of VGCCs: L, N, P/Q, R, and T-types (*12, 13*). We utilized pharmacological antagonists to inhibit these channels selectively. This approach unveiled that L- and N-type Ca^++^ channels are involved at the earliest stages of the shrinkage following axotomy. Their respective blockers, nifedipine (NIF) or ω-conotoxin GVIA (CNX), effectively inhibited cell size reduction after axotomy (Fig. 2A). NIF-mediated inhibition was most effective in the first 5 min while CNX exerted a more consistent inhibition. Blocking R-type channels with SNX-482 (SNX) effectively prevented shrinking after 15-60 mins post axotomy. Inhibition of P/Q type channels with ω-agatoxin-IVA (AGX) and T-type Ca^++^ channels with mibefradil (MIB) proved ineffective. The results of a prior work depolarizing rat DRG neurons with high K^+^ and examining the contribution of VGCCs to Ca^++^ influx are consistent with the preponderance of L- and N-type channels in the early stages of shrinkage and the major contribution of R-type channels later on. (*12*). VGCCs are classified as either high or low-voltage-activated types. The level of their expression and expression ratio among subtypes of primary sensory neurons differ; these are subject to change in culture (*13–15*). Our findings demonstrate that Ca^++^ influx through the high-voltage-activated L-, N-, and R-type channels induce shrinkage in cultured DRG neurons. Confirmation of these results *in vivo* remains to be investigated.

**Fig. 2.**
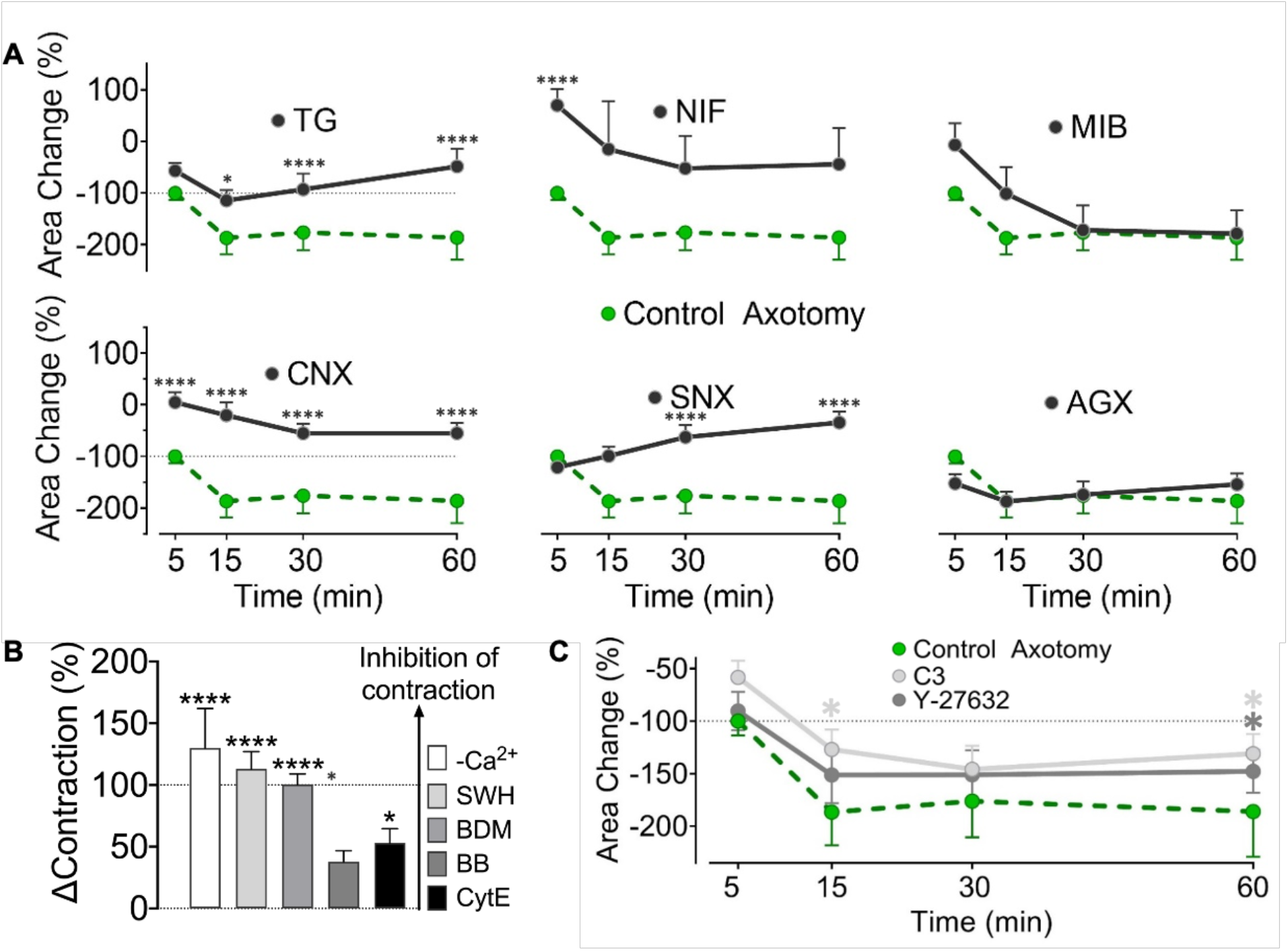
Mechanisms of calcium elevation and AMC after axotomy. (**A**) Effect of inhibition of voltagegated calcium channels (VGCC) and internal Ca^++^ stores on surface area changes after axotomy. VGCC subtypes L, N, P/Q, R, and T, were blocked with nifedipine (NIF) (n=4/46), ω-conotoxin GVIA (CNX) (n=3/30), ω-agatoxin-IVA(AGX) (n=3/23), SNX-482(SNX) (n=3/18) and mibefradil (MIB) (n=3/28) respectively. Thapsigargin (TG) (n=4/44) was used to deplete internal Ca^++^ stores before axotomy. **(B)** Effect of inhibition of AMC on surface area change 5 mins after axotomy in Ca^++^-free medium (n=3/38), myosin inhibitors butanedione monoxime (BDM (n=3/30)) and blebbistatin (BB) (n=4/48) and actin inhibitors cytochalasin E (CytE) (n=3/33) and swinholide A (SWH) (n=3/41). The data are normalized relative to control axotomy (baseline) and presented as mean ± SEM for easier interpretation; the taller the bar, the more potent the inhibition of shrinkage. **(C)** Effect of inhibition of RhoA signaling with clostridium botulinum exoenzyme C3 (n=3/20) or RhoA’s effector ROCK with Y-27632 (n=3/30) on surface area change after axotomy. Mann-Whitney U test was used to compare surface area changes in each group to the changes 5 mins (B) or at indicated time points (A, D) after control axotomy (n_5_=6/65, n_15_=6/28, n_30_=6/43, n_60_=6/60). Error bars: SEM; n=number of animals used/number of cells analyzed; for all tests: *p<0.05, ****p<0.00001.

Reports show that axotomy can mobilize internal Ca^++^ stores (*16*). To understand the contribution of endoplasmic reticulum stored Ca^++^ to cell shrinkage, we used thapsigargin (TG), a Ca^++^ ATPase inhibitor, in a two-stage experiment described elsewhere (*17*). First, neurons were incubated with TG to empty the internal Ca^++^ stores. Then, we performed axotomies and analyzed shrinkage to test the effect of extracellular Ca^++^ alone. Eliminating internal Ca^++^ release did not alter the shrinkage during the first 5 min following axotomy (Fig. 1A). However, cell shrinkage could not continue for more than 15 minutes without internal Ca^++^ reserves.

In this study section, we demonstrated that extracellular Ca^++^ coming through VGCCs initiates axotomy-induced shrinking, which is later maintained by Ca^++^ levels from internal stores. The interplay of multiple mechanisms probably determines the net increase in cytosolic Ca^++^. In this sense, the potential role of store-operated Ca^++^ entry (SOCE), typically induced by depletion of internal stores, may also be considered (*18*). However, we obtained indirect evidence that SOCE is unlikely to share responsibility for axotomy-induced shrinkage. According to previous reports, SOCE is activated within one hour of the TG application(*19*). In our experiments, the time from TG application to axotomy was 90 minutes, enough to induce SOCE. If SOCE played a significant role in the axotomy-induced increase in cytosolic Ca^++^, it is expectable that shrinking would continue for 60 minutes after the axotomy. Therefore, we indirectly conclude that SOCE may not be essential for axotomy-induced shrinkage.

The following hypothesis was that the axotomy-induced elevation in cytosolic Ca^++^ may lead to shrinkage through active contraction and the cytoskeletal reorganization necessary for size change. To provide evidence, we first conducted *in vitro* axotomies in Ca^++^ - free medium or in the presence of inhibitors of actin (cytochalasin E -CytE and swinholide A - SWH) or myosin (blebbistatin – BB and 2,3-Butanedione monoxime-BDM). Lack of extracellular Ca^++^ or addition of SWH, CytE, or BDM significantly inhibited neuronal shrinkage while BB was ineffective (Fig. 2B). CytE prevents polymerization of actin while SWH severs f-actins (*20, 21*). Expectedly, the effect of CytE was substantially less than SWH, indicating that the pre-existing f-actins may still mediate part of the contraction despite the elimination of new actin polymers. Failure of BB to counteract the shrinkage may be due to its high selectivity against myosin subtypes (*22*). However, BDM, a more general myosin inhibitor that blocks phosphorylation of myosin light chain (MLC) kinase (*23*), effectively blocked cell-shrinkage.

### RhoA signaling is not essential but contributes to the AMC

To understand how AMC is initiated, we next focused on the Rho GTPases and their downstream effectors, Rho-associated kinases (ROCKs), as potential signaling mechanisms. In non-muscle cells, such as neurons, RhoA and ROCK govern actomyosin contractility in a Ca^++^-dependent manner and moreover they are activated by axotomy (*24, 25*). We performed axotomy in the presence of RhoA inhibitor exoenzyme C3 and ROCK inhibitor Y-27632. RhoA inhibition significantly reduced shrinkage at the 15^th^ and 60^th^ min post axotomy, while ROCK inhibition decreased cell shrinkage only at minute 60 (Fig. 2C). Y-27632 displayed marginal effects, possibly because ROCK enhances AMC only through MLC by inactivating its inhibitor MLC phosphorylase (*26*). However, RhoA has an additional effector, diaphanous-related formin 1 (mDia1), which activates profilin, an actin-polymerizing and stabilizing factor (*27*). Our data indicate that RhoA activation contributes to AMC minutes after axotomy and is only partially responsible for the shrinkage. In a relevant study, an endothelial cell stretch model demonstrated a similar temporal relationship with Ca^++^ increase, RhoA activation, and AMC(*28*). We suggest that axotomy-induced membrane depolarization is the first step in the likely mechanism of AMC. Next, Ca^++^ influx directs the contraction while activating the RhoA system, which may aid in boosting actin polymerization and disinhibiting MLC.

### Calpains digest the cytoskeleton to enable cell shrinkage

Beneath the plasma membrane lies a contractable actomyosin cortex that participates in various physiological processes, including alteration of cell shape and the control of intracellular hydrostatic pressure (*29*). To make changes in a cell’s size or shape, the tension created by AMC must overcome the opposing forces exerted by relatively static structural elements of the cytoskeleton. Therefore, we reasoned that axotomy might cause the activation of proteolytic enzymes, namely calpains, which could break down cytoskeletal proteins and reduce the opposing forces. To test this hypothesis, we axotomized cultured neurons in the presence of a general calpain inhibitor PD150606 (PD). We found that this approach almost completely prevented cell shrinkage (Fig. 3A). Calpains are activated by increased intracellular Ca^++^, targeting many functional and structural proteins, which make them accountable for various pathological changes in neurodegenerative processes (*30, 31*). Although identifying calpain substrates is out of the scope of this study, we suspected that it might be spectrin because it forms a pre-tensed supporting mesh under the plasma membrane and tubulin, which constitutes the major solid network opposing the compression forces (*30, 32*). On the other hand, reports suggest that actins and myosins are not the primary targets of calpains (*33*). Thus, we presumed that calpain inhibition did not preclude AMC, but neurons could not shrink due to the resistance of the intact cytoskeleton. Indeed, when calpains and AMC were blocked simultaneously, neurons’ response to axotomy was reversed to swelling instead of shrinking. In the earlier experiments, BDM alone prevented shrinkage but did not cause swelling (Fig. 1B). How simultaneous inhibition of calpains led to the swelling is unclear. Yet, it may indicate that calpains’ actions after axotomy are possibly as diverse as their many substrates (*33*).

**Fig. 3.**
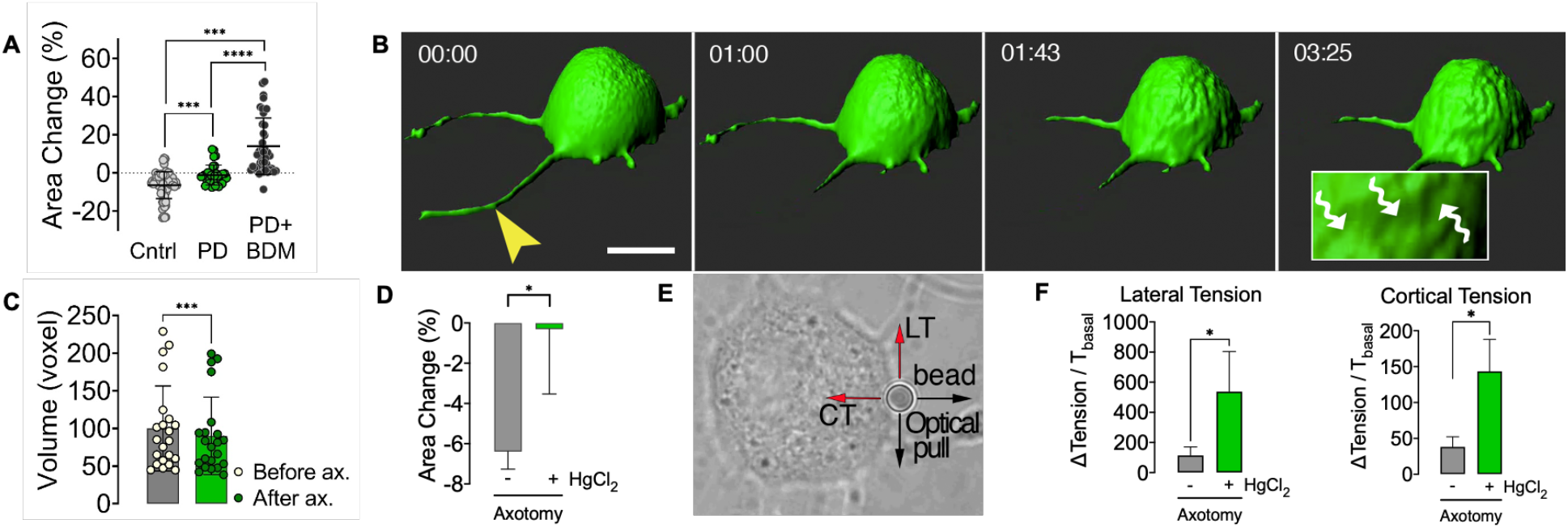
Characteristics of axotomy–induced shrinkage. **(A**) Scatter dot blot shows surface area change 5 mins after axotomy in the presence of calpain inhibitor PD150606 (PD) alone (n=3/33), in combination with the myosin inhibitor butanedione monoxime (BDM) (dots, n=4/41) and in control conditions (n=6/65). Kruskal Wallis (A) and Mann-Whitney U-tests were used for comparisons. **(B)** Representative threedimensional confocal images show an axotomized neuron over time. Wavy arrows in the inlet point to depressions on the cell surface due to shrinkage (scale bar, 15 μm; arrowhead points to the site of axotomy). **(C)** Mean volume of DRG neurons before and 5 mins after axotomy (n=3/22). Wilcoxon Signed Ranks Test was used for comparison. **(D)** Comparison of surface area change 5 mins after axotomy with AQP inhibition (n=3/20) to the control axotomy (n=6/65). **(E)** A Representative bright-field image shows the measurement of membrane tensions by pulling an attached polystyrene bead with an optical tweezer. (**F)** Changes in the membrane tensions (T) due to axotomy (n=4/48) and the effect of aquaporin (AQP) channel inhibition with HgCl_2_ (n=3/20). Mann-Whitney U test was used for comparisons in D and F. Error bars: SEM; n=number of animals used/number of cells analyzed; for all tests: *p<0.05, ***p<0.0001, ****p<0.00001.

### Isotonic contraction pumps water out of the axotomized neuron

We next sought to understand whether the change in cell surface area after axotomy means a change in volume as the opposite is also possible since neurons may get elongated vertically while shrinking laterally. We cultured DRG neurons from EGFP mice and acquired three-dimensional images with a fast confocal microscope during axotomy. We determined that neuronal volumes were significantly reduced concordant with the change in surface area (Fig. 3B, C, and movie S5). To explain the volume loss, we proposed that, by contraction, water may be extruded from the cells through aquaporin channels (AQP) (*34*). Indeed, when we blocked AQPs with HgCl_2_, neuronal size did not change upon axotomy (Fig. 3D). To further characterize the contraction, we measured the plasma membrane tensions of neurons before and after axotomy with a special microscope combining laser microdissection and an optical tweezer (Fig. 3E). The slight increases in lateral and cortical tensions caused by axotomy were significantly augmented by inhibition of AQPs with HgCl_2_ (Fig. 3F). This observations strongly suggests that axotomy causes an isotonic type of contraction characterized by a decreasing cell volume and only slightly elevated membrane tensions consistent with the Law of Laplace (*35*). Increased membrane tensions show that cellular contraction becomes isometric in nature when volume change is inhibited (Fig. 3D, F).

### Shrinkage increases the survival chance of axotomized neurons *in vitro* and *in vivo*

Our results thus far provide evidence that neurons respond to axotomy by shrinking under the influence of AMC and calpain activity, which forces water out of the cell through AQP channels. Thus, we hypothesized that shrinkage could help neurons survive by preventing swelling. We ran several *in vitro* and *in vivo* experiments to test this hypothesis.

In culture, when the contraction was inhibited by BDM, more neurons died progressively over 24 hrs incubation period (Fig. 4A and supplementary text). The surviving axotomized neurons for the next 1 and 24 hrs were those that had shrunk, and those that died had not shrunk by the 5^th^ minute post axotomy (Fig. 4B). The chance of survival after axotomy was correlated to the degree and persistence of shrinkage (fig. S1 and supplementary text). Using *in vivo* models, we sought to confirm the biological relevance of these *in vitro* findings suggesting that shrinkage favors neuronal survival. Using the same technique utilized for *in vivo* axotomy (*11*), we constructed an imaging window over L2 DRG of Tau - EGFP mice whose neuronal soma was visibly labeled (*36*). The basic experimental paradigm was to cut the peripheral nerve emanating from the DRG, thus axotomizing all neurons. Propidium iodide (PI) was injected intravenously to monitor cell death. In some experiments, BDM was intrathecally administered before nerve cut. With a two-photon microscope, we took images before, at 5 minutes, and 24 hrs after axotomy. The nerve cut caused an apparent reduction in size, which was even more significant after 24 hrs (fig S2). BDM prevented neuronal shrinkage and significantly increased neuronal death compared to control animals, while the sham group with only BDM injection showed no neuron loss (Fig. 4C, D and fig S2). As demonstrated by *in vitro* experiments, surviving neurons were those that had shrunk. Expectedly, DRG neurons from Thy1-GFP animals pre-injected with BDM did not shrink upon single-cell axotomy *in vivo* (fig S2).

**Fig. 4.**
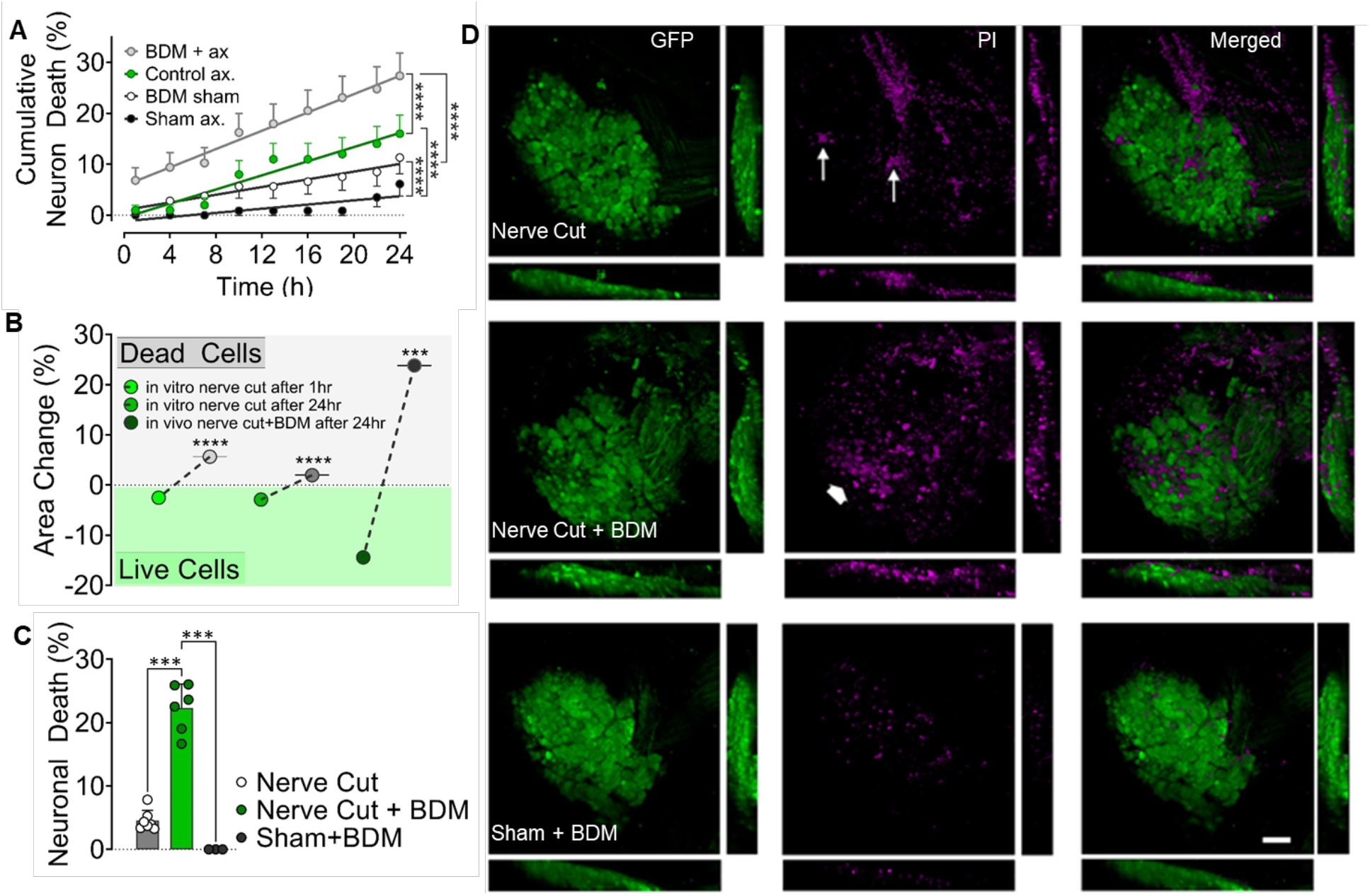
Relationship of axotomy-induced contraction with neuronal survival. **(A)** Cumulative death rates of axotomized and control neurons during 24hrs incubation period with and without inhibition of AMC by butanedione monoxime (BDM) (n=3/117 (BDM+ax), 3/100 (Control ax), 3/106 (BDM sham), 3/114 (Sham). Groups were compared using a Repeated - measures Mixed Model. **(B)** Surface area changes of neurons that survived and died after *in vitro* axotomy (n=16/303 (live 1 hr), 16/113 (dead 1 hr), 16/160 (live 24 hr), 16/266 (dead 24 hr)) and *in vivo* nerve cut (n=6/11 (live), 6/10 (dead)). Note that *in vitro* data were pooled from various experimental groups, including those with pharmacological agents. Mann-Whitney U test was used for comparisons. **(C)** Rate of cell death in DRGs 24hrs after peripheral nerve cut or sham operation and effect of inhibition of AMC with BDM *in vivo*. A comparison of neuronal death rates was performed with Kruskal Wallis and Mann-Whitney U tests, and the data are shown as mean ± SEM (n=7 (nerve cut), 6 (nerve cut+BDM), 4 (sham + BDM)). **(D)** Representative confocal images of DRGs from *in vivo* experiments in C. Arrows point to a few PI-stained dead neurons, while the thick arrow shows a large group of them (scale bar, 100 μm). Error bars: SEM; n=number of animals used/number of cells analyzed; for all tests: ***p<0.0001, ****p<0.00001.

An interesting observation we made but not quantitatively analyzed in this study was that contraction of soma could constrict and amputate the proximal stump when axotomy is very close, which represents a fast and efficient way of membrane resealing, potentially increasing the chance of survival when the risk of death is high (movie S6). In line with our observations, Kelley et al. demonstrated in an *in vivo* model that traumatic axonal injury close to neuronal soma in rat thalamus results in secondary axotomy within 15 min and is accompanied by an efficient resealing (*37*).

## Conclusion

This study describes an axotomy-induced AMC coupled with proteolysis that shrinks injured neurons to pump out water and prevent deadly swelling (Fig. 5). The postulated pathway begins with axotomy-induced membrane depolarization, opening high-voltage-activated Ca^++^ channels. AMC and calpain enzymes are activated by elevated cytoplasmic Ca^++^ (internal Ca^++^ stores can contribute) and break down cytoskeletal proteins to lessen the soma’s ability to resist shrinking. The cell contraction is partially sustained by RhoA signaling and occurs isotonically pumping water out of the cell through AQP channels.

**Fig. 5.**
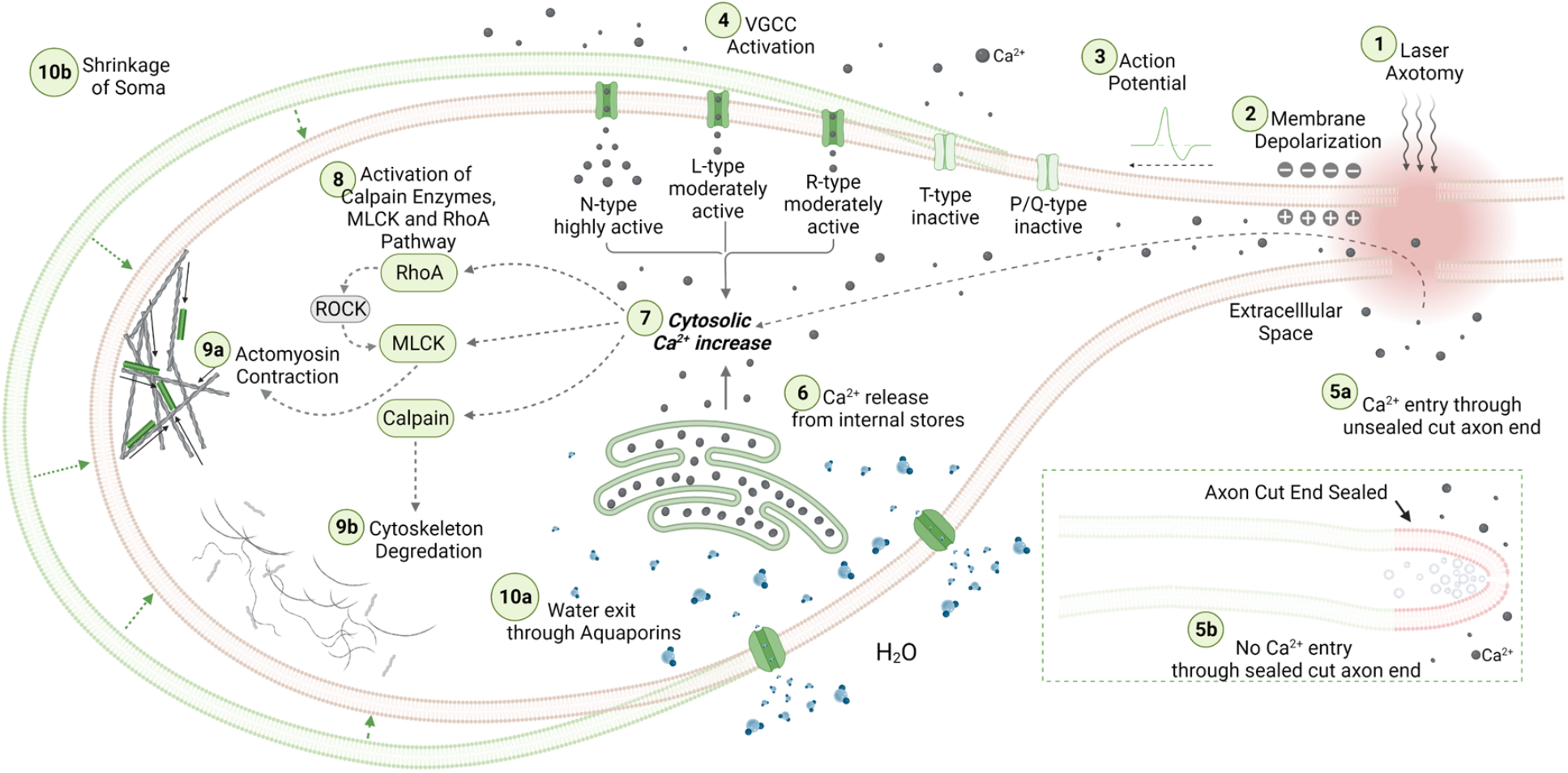
Schematical overview of the involved mechanisms in cell shrinkage upon axotomy: See main text for a detailed description.

The tell-tale sign of this protective response is a reduction in somatic size both in central and peripheral neurons.Various other studies have reported that neurons’ diameter decreases after axonal injury, which was often intrepretted as atrophy (*38–43*). Although atrophy is a likely outcome of an axotomy in the long term, shrinkage in cholinergic septal neurons as early as 24 hrs and retinal ganglion neurons three days after axotomy may not technically be called atrophy (*39, 42*). In the existing literature, the contractile capacity of neurons has been mainly associated with neurites and growth cones (*9, 44*). An exception relevant to our findings was a report by Wan et al. back in 1995, stating that “*a plasma membrane-linked contractile machinery (presumably actomyosin) might contribute to the neurons’ mechano-osmotic robustness by restricting water influx”* during hyposmotic challenge (*45*). We have demonstrated that water leaves the shrinking neurons via AQP channels. AQP1 is expressed in DRG neurons and upregulated after nerve injury (*46*), and AQP2 expression is induced after nerve constriction (*47*). HgCl_2_, by which we blocked water efflux, is primarily an AQP1 inhibitor but not strictly selective (*48*). Therefore, we do not exactly know which subtypes of AQPs are responsible for water efflux. We investigated the mechanism of shrinkage using mainly DRG neurons. Although we did not conduct a differential analysis, heterogeneity among different subtypes of these neurons in shrinkage response to axotomy can be expected due to varying constitutional expression of molecular components of the event. This heterogeneity is even more likely in central neurons, where we observed minor average size reduction with more significant deviations across individual cells. On the other hand, our observation that the initial size of DRG neurons, which roughly defines different subpopulations (*49*), does not affect the unique consequence of the shrinkage - the increased chance of survival-suggests that phenotypic differences do not preclude the significance of the phenomenon. We believe that the interaction of shrinking neurons with neighbouring cells may have significant consequences, such as mechanical signaling (*50*).

In this study, we have revealed a critical phenomenon that protects neurons after axotomy. This mechanism may have far-reaching implications in various pathological conditions involving the nervous system, from direct traumas to neurodegenerative diseases associated with primary or secondary axonal injury.

## Materials and Methods

### Animals

The use of animals in this study was approved by Istanbul Medipol University Animal Experimentation Ethical Committee. Maximum care was paid with total commitment to the 4R principles to use the minimum number of animals necessary for the research aims and to minimize suffering throughout the study. All procedures were conducted in compliance with European Council Directive 2010/63/EU.

Breeding colonies of all mice strains were obtained from The Jackson Laboratory and bred at Istanbul Medipol University Experimental Animal Facility. These were general GFP - B6 ACTb-EGFP (stock no: 003291) (*51*), TauEGFP knock-in (stock no: 004779) (*36*), Thy1-GFP line M (stock no: 007788) (*10*), cross breeds of cre-dependent GCaMP6s - Ai96(RCL-GCaMP6s) (*52*) (stock no: 024106) and Vglut2-ires-cre (stock no: 016963) (*53*) and non-transgenic BALBc (stock no: 000651)mice.

### Neuron cultures

#### Adult primary sensory neuron cultures

Dorsal root ganglia (DRG) were removed from adult mice, and primary sensory neurons were isolated as described before (*54*). Briefly, animals were anesthetized with an I.P. injection of ketamin (Pfizer) and sacrificed by cervical transection. The vertebral column was cut out and quickly transferred into a dissection plate with RPMI1640 medium (Sigma). DRGs were collected under a stereo microscope and digested by collagenase (Sigma, 100 U/ml, 50 min) and trypsin (Sigma, 1mg/ml, 15 min) treatments, after which they were triturated through pipet tips of narrowing bores for 15 minutes and finally through a 26-gauge injector needle. The cell suspension obtained was treated with DNAse, spun at 120 g for 3 min, and the pellet was resuspended in Neurobasal A supplemented with B27 (Thermo Fisher), glutamine (Glutamax-Thermo Fisher), and antibiotic solution (Sigma) (NBA-B27). To eliminate nonneuronal cells, a gradient centrifuge technique was used. The cell suspension was carefully pipetted on top of a three-layer percol (Sigma) gradient (60%, 35%, and 10% from bottom to top) prepared in NBA-B27 and spun at 3000 g for 20 min in a centrifuge at 4°C. The neurons collected from 35% layer were washed with NBA-B27 and spun once more at 120 g for 3 min, after which the supernatant was discarded, and the pellet was resuspended in NBA-B27.The suspension was transferred on 35 mm diameter glass-bottomed Petri dishes (WPI) coated with poly-L-lysine and laminin and incubated at 37°C with 5% CO_2_.

#### Newborn hippocampal neuron cultures

Postnatal (p1-p3) mice were sacrificed with CO_2_, and brains were collected under aseptic conditions. Hippocampi were surgically removed in a dissection medium (L15, 1% antibiotic-antimycotic solution), cut into small pieces, and digested with papain (Sigma, 12.5 U/ml, 45 min) with agitation at 4 °C. Afterward, the suspension was triturated via serial pipetting, and papain was inhibited with 10 % FBS containing dissection medium for 15 min at 4 °C. Then, cells were pelleted by centrifugation at 180 g for 5 min at 4°C, the dissection medium was discarded, and cells were plated on poly-D-lysine coated glass bottom culture dishes and incubated in NBA-B27 at 37°C with 5% CO_2_.

### Microscopic imaging and in vitro manipulations

For microscopic imaging, the following systems (all from Zeiss) were used: LSM780 NLO, LSM880, and LSM7 MP laser scanning confocal and two-photon microscopes, Cell Observer time-lapse imaging system, and Palm Combi system. Various modules of Zeiss Axiovision and Zen software were used for time-lapse imaging of multiple positions at desired intervals, threedimensional scanning, and Ca^++^ signal imaging. Axotomy was performed with the femtosecond infrared lasers of two-photon microscopes in vivo, and UV lasers fitted to LSM880 and Palm laser microdissection - optical tweezer combi system in vitro.

#### In vitro axotomy

A brief pulse (0.5-1 sec) of UV (337nm or 355nm) laser beam was focused on axons about 50-100 μm distal to the soma emitting approximately 1–30 pulses per second, each lasting 3 nsec and delivering approximately 300 μJ of energy. For sham operations, an empty spot 50 μm away from the cell body was irradiated similarly.

#### Membrane tension measurement

The optical tweezer controlled by Palm software with a force measurement module was used for membrane tension measurements. For this, poly-L-lysin-coated 3 μm-diameter polystyrene beads (Sigma) were carried and attached to the neuronal plasma membrane by the optical trap. Then they were pulled vertical or parallel to the plasma membrane with the optical trap to measure cortical and lateral membrane tensions, respectively. The force resisting the pull was measured in piconewtons and reported by the software.

### In vivo experiments

#### Vertebral window surgery

Construction of an imaging window over lumbar DRGs was performed according to the method described by Chen et al. (*11*). Briefly, the mice were anesthetized with 2% isoflurane (30% O_2_), and a small incision was made in the dorsal skin at the level of L1-L3 of the spine, paraspinal muscles and ligaments connected to right transverse processes of L1 to L3 were carefully dissected using fine scissors to expose DRG. The mouse was then placed lying on its left side onto a custom-made vertebral holder, which allows access to L2 DRG for in vivo microscopy. Carefully removing surrounding muscles and connective tissue around the L2 DRG using a fine tweezer, DRG was exposed. To achieve optical clearance and stabilization, DRG and surrounding bone structures were covered with silicon adhesive Kwik-Sil® (World Precision Instruments), and sealed with 3-mm diameter round cover glass. The cover glass was fixed to vertebrae using cyanoacrylate glue and dental cement.

#### Peripheral nerve cut

TauEGFP knock-in mice were used to perform axotomy in all neurons within the DRG by cutting the peripheral nerve of DRG. This strain was preferred for these experiments due to strong expression of EGFP in neuronal soma, which enabled deep imaging and analyzing surface area changes. Using the two-photon microscope (LSM 7MP), neurons in L2 DRG were imaged through the vertebral window with 10x/0.45NA air objective, and 820nm excitation laser before and after its peripheral nerve was completely cut with surgical scissors.

#### In vivo axotomy

Thy1-GFP line M mice were preferred for single cell axotomy as seldom labeled DRG neurons with clearly visible axons were easier to identify and analyze after the procedure. Single-cell axotomy was performed by irradiating axons with Ti:Sa laser about 100 μm distal to the cell body. For this purpose, the laser was adjusted for a single spot scan with 100 iterations at 820 nm, delivering ~270mJ power at the focal plane.

#### Myosin inhibition and analysis of neuronal survival

Intrathecal delivery of myosin inhibitor 2,3-Butanedione monoxime (BDM) was performed as previously described (*55*). Briefly, a Hamilton syringe (Hamilton 80314, Hamilton, Reno, NV) with a 32-gauge needle was inserted between the L2-L3 vertebrae to the subarachnoid space. Access to the intrathecal space was confirmed by reflux of cerebrospinal fluid. Five microliters of SF containing 50mg/ml BDM was injected intrathecally 30 minutes prior to in vivo experiments. To determine the cell death in DRG neurons, 50 μl of 1/1000 propidium iodide (PI, Molecular Probes), which stains nuclei of dead cells, was injected intravenously via the tail vein 1 hr before the microscopic imaging.

### Patch-clamp recordings

Patch-clamp pipettes were made of borosilicate glass (Sutter Instrument Borosilicate Glass with flament. O.D.: 1.5 mm, I.D.: 1.17 mm 10 cm Length, Novato, CA, USA). The standard extracellular bath solution contained (in mM): 130 NaCl, 2 CaCl_2_, 10 HEPES, and 13 D-glucose, pH adjusted with NaOH to 7.35. The osmolarity of the solution was 270 mOsmol/L. The pipette (internal) solution for patch-clamp recording contained (in mM): 130 K-gluconate, 5 EGTA, and 10 HEPES (270 mOsmol/L; pH 7.2) (adjusted with NaOH). The pipette resistances were about 4 – 6 MΩ. Also, the neurons were held at a potential of −70 mV. The resistance was made to be gigaseal (2 - 10 GΩ), and then the whole cell configuration was started with negative pressure. The Whole-cell patch clamp technique was used to perform current-clamp recording by using an amplifier (Axon CNS MultiClamp 700B). Low-pass-filtering at 5–10 kHz was used during data recording. Sampling rates for current and voltage records were 10–20 kHz. A Digidata 1550B interface (Axon Instruments, Foster City, CA, USA) was used for the digitization of data, and a computer was used for storage and further analysis. pClamp software (version 11; Axon Instruments) was used for the generation of stimulus, acquisition, and offline analysis of data.

To record from neurons during axotomy, the patch clamp rig was mounted on a Palm laser microdissection microscope, and after a giga-seal was created, a selected axon of the patched neuron was cut with a laser beam as described earlier.

### Calcium imaging

Cultured DRG neurons of GCaMP6s - Ai96 mice were imaged with a confocal microscope before, during, and immediately, then every 5 mins after axotomy. Change in green fluorescence emission was analyzed using Zen software.

### Use of blocking reagents in vitro

Neurons were incubated with the blockers for one hour before microscopic manipulations except for exoenzyme C3, whose incubation period was 6 hrs. All inhibitors were purchased from Sigma, except exoenzyme C3, which was obtained from Abcam.Cytochalasin E (CytE, 200 ng/ml) (*21*) and swinholide A (SWH, 500 ng/ml) (*56*) were used to depolymerize actin. Myosin activity was blocked with blebbistatin (BB, 100 μM) (*57*) and 2,3 butanedione monoxime (BDM, 50mM) (*23*). Voltage-gated calcium channel subtypes L, N, P/Q, R, and T were blocked with nifedipine (NIF, 10μM) (*58*), ω-conotoxin GVIA (CNX, 1μM) (*59*), ω -agatoxin-IVA(AGX, 200nM) (*60*), SNX-482(SNX, 200nM) (*60*) and mibefradil (MIB, 10μM) (*61*), respectively. To test the contribution of internally stored Ca^++^ to the shrinkage, neurons were incubated with thapsigargin for 60 min (TG, 2μM) (*17, 62*) to empty internal Ca^++^ stores. Since store-operated Ca^++^ entry is induced by depletion of internal stores, these experiments were performed in Ca^++^ - free DMEM (*17*). Then the medium was replaced with NBA-B27, and 30 min later, axotomies were performed.

PD150606 (50μM) (*63*) was used as a calpain inhibitor. Aquaporin channels were blocked using HgCl_2_ (100μM) (*64*). Ras homolog gene family member A (RhoA) and its downstream effector Rho-associated protein kinase (ROCK) were inhibited with clostridium botulinum exoenzyme C3 protein (C3, 500 nM) (*65*) and Y-27632 (10 μM) (*66*), respectively.

### Image Analyzes

Changes in surface area and volume of neurons were analyzed using ImageJ software. Fluorescence intensity measurements were performed with Zeiss Zen software.

### Statistical analysis

IBM SPSS V24 and SAS V9.4 were used. The distribution of the data was evaluated with Kolmogorov -Smirnov test. Though some data sets had a normal distribution, most of them did not, therefore, non-parametric tests were used for group comparisons. The distribution of a continuous marker was compared among two or more levels of a factor of interest by using Wilcoxon-Mann-Whitney or Kruskal-Wallis tests. Change of marker overtime was modeled through random-coefficient models using the MIXED procedure in SAS, where spatial-power of time-lag covariance structure was used to account for the correlation structure among measurements. Similarly, repeated-measures models with variance-components covariance structure were also constructed using the MIXED procedure. To estimate the impact of a given predictor on the likelihood of survival, univariable Logistic regression was used and the significance of the predictor was assessed through both p-values and Area Under the curve (AUC).

## Author contributions

Conceptualization: GÖ, MŞA; Methodology: GÖ, AH, MK, EE; Investigation: GÖ, MŞA, SB, ENY, CÖ, EYK, EKO, NA, NC, EE (1), AH; Visualization: GÖ, MŞA, ENY, EE (1), EE (2); Funding acquisition: GÖ; Project administration and supervision: GÖ; Writing – original draft: GÖ, Writing – review & editing: GÖ, MK, EE (2).

## Acknowledgments

This research was funded by Yüzüncü Yıl University Directorate of Scientific Research Projects, grant TF073 (GÖ) and the Scientific and Technological Research Council of Turkey (TÜBİTAK), grant 107S358 (GÖ).

## Data and materials availability

All data will be deposited to an online repository upon acceptance of the manuscript for publication.

