## supplementary text for "Active shrinkage protects neurons following axonal transection"

### Neurons protect themselves by active shrinkage upon axonal transection

#### Supplementary Text

Membrane Depolarization by Axotomy

Axotomy caused about 16 mV depolarization on average and created 1-321 action potentials in 9 of 16 neurons. The firing rates of action potentials were 1.4-26 Hz, which lasted up to 37 secs and membrane stayed depolarized during the recording period of 25 mins. Resting membrane potential 24 hrs after axotomy had recovered to control conditions (-51.1 mV(control) vs. -57.1 mV (axotomized), p>0,05); but when re-axotomized only 1 of 12 neurons generated action potential (p=0.016) while all got depolarized with a voltage change similar to first axotomy.

Shrinkage – Survival Relationship

Cumulative cell-deaths were reported for each hour from baseline to 24 hour for each group (Fig. 4). As this is an example of repeated measures data and the independent assumptions among observations may be violated, we employed mixed-modelling approach considering the repeated measures structure, using the MIXED procedure in SAS Version 9.4, where cumulative cell death was the outcome variable with time and experimental group and their interactions being the predictors. Time*experimental group interaction was found to be highly significant (p<0.0001). The slope estimate for the sham group was 0.19 (p<0.0001), suggesting one death in about every 5 hours. Slope of BDM-Sham group was about 0.19 unit higher compared to that of the Sham group (p=0.0004, slope estimate=0.38), suggesting that BDM-Sham group had one more additional cell death for about every 5 hours compared to the sham group. Axotomy group had a slope estimate that was 0.53 unit higher (p<0.0001, slope estimate=0.71) than that of the Sham group, resulting in about one more additional cell death in every two hours. BDM-Axotomy group had about 0.85 excess cell-death per hour compared to Sham group (p<0.0001, slope estimate=1.04) and one more additional cell death for about every 3 hours compared to the control axotomy (p<0,0001). The analyses of pooled data derived from images taken at the 5, 15, 30, 60$^{th}$ minutes and 24hrs after axotomy from in vitro groups revealed following results: 1. According to the univariable logistic regression model we developed, every 1% shrinkage by 5$^{th}$, 15$^{th}$, 30$^{th}$ and 60$^{th}$ minutes after axotomy resulted in 9, 7, 4 and 3% increase in likelihood of survival, respectively (p<0,0001 for each time points). 2. Cut off points for 5, 15, 30 and 60 minutes after axotomy were 0%, -8%, -3% and -2% size change respectively; below which the chance of survival started to become significantly higher (fig. S1). We also performed a longitudinal analysis of the data and constructed random-coefficients models where each cell had its own random intercept and slope to describe its profile overtime. In these models, initial cell size, which roughly defines different phenotypic subpopulation of DRG neurons, was not significantly associated with survival (p=0.58) and it had no time interaction. However, survival status and time had a significant interaction as shown in the fig. S1B, suggesting that the neurons that were able to continuously shrink were more likely to survive.

**Supplementary Figures**

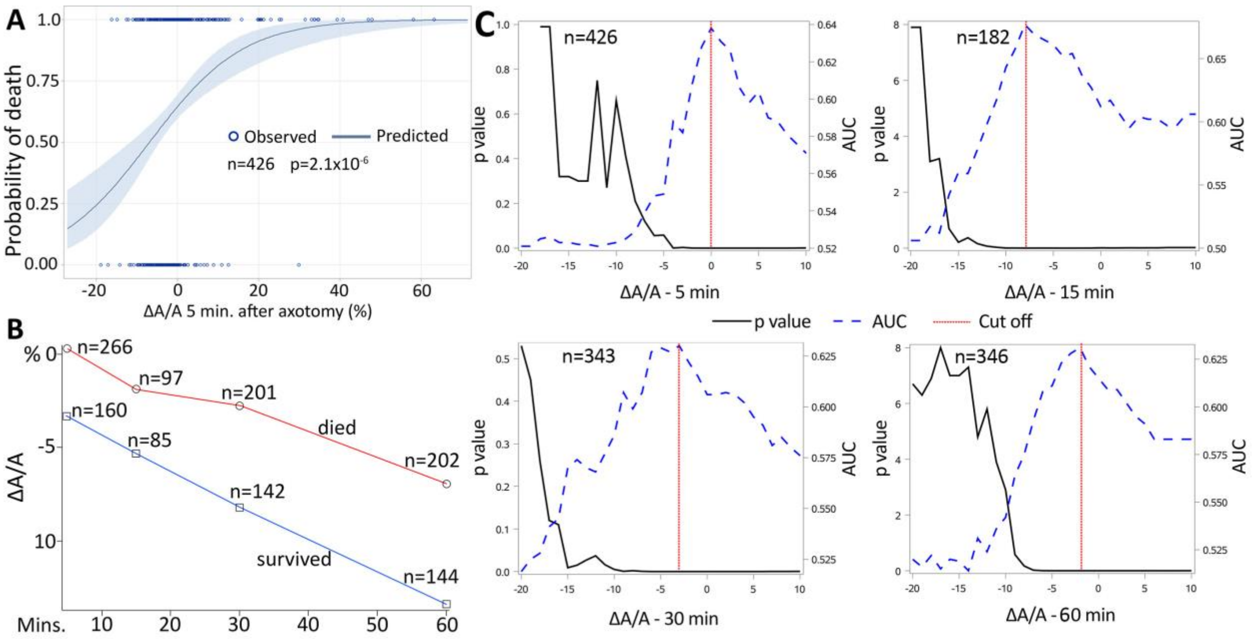


**Fig. S1. Correlation of shrinkage with survival**. **(A)** Univariable regression model showing correlation between degree of shrinkage and probability of neuronal death. **(B)** Longitudinal model of shrinkage of neurons that survived and died after axotomy. (C) Cut off values for survival – shrinkage relationship at 5, 15, 30 and 60 min following axotomy. Data from multiple experimental groups were pooled.

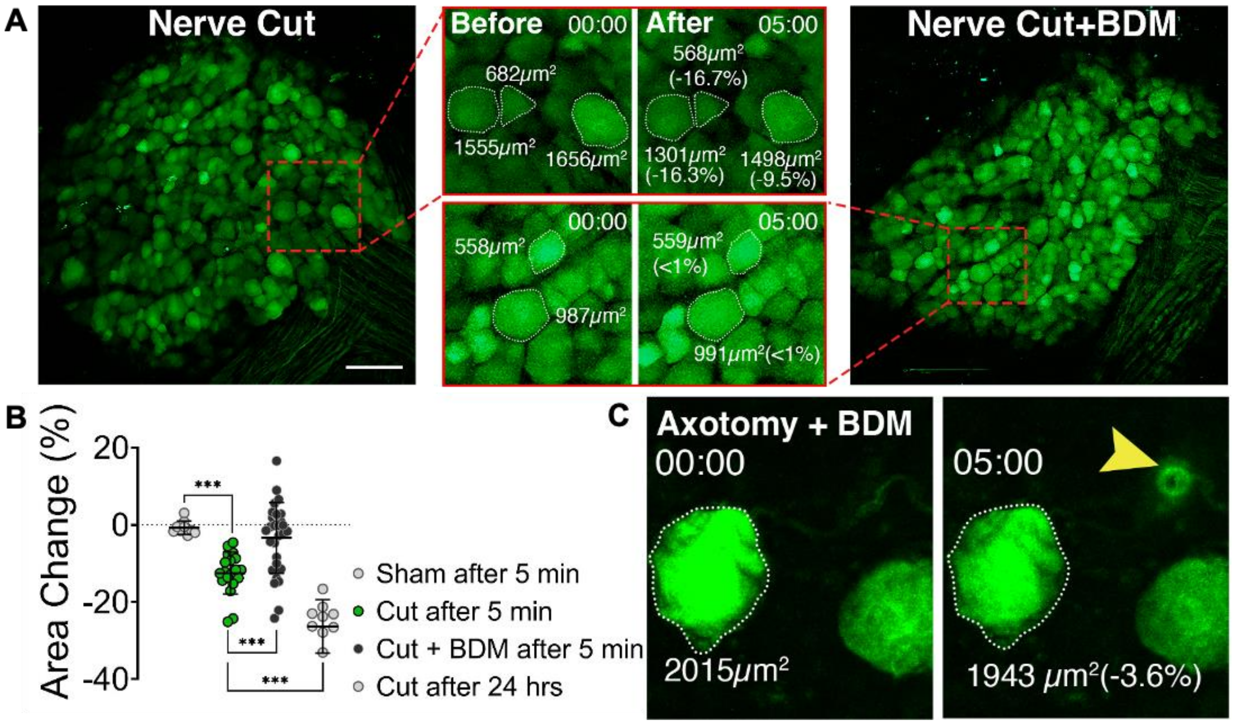


**Fig. S2. Shrinkage in DRG neurons after in vivo nerve cut and single neuron axotomy with two-photon microscope. (A)** Representative images of DRGs from TauEGFP mice before and after peripheral nerve cut and effect of inhibition of actomyosin contraction with myosin inhibitor butanedione monoxime (BDM) (scale bar, 100 μm). **(B)** Scattered dot plot shows surface area changes 5 min and 24 hrs after intervention in control and nerve cut groups and effect of BDM. Mann-Whitney U test was used for comparisons; (n=7/19 (cut 5 min), 7/10 (cut 24 hrs), 6/29 (cut+BDM 5 min), 3/9 (sham 5 min)), ****$p<0.00001$. **(C)** Images of a neuron from Thy1-GFP mouse axotomized *in vivo* in the presence of BDM, arrowhead points to the site of injury (compare it to Fig. 1A).

#### Movie Legends

**Movie S1.**

Axotomy and shrinkage of a cultured DRG neuron.

**Movie S2.**

$Ca^{++}$ increase by axotomy in a DRG neuron expressing $Ca^{++}$ sensor - GCaMP6s.

**Movie S3.**

$Ca^{++}$ entry into a DRG neuron after axotomy through unsealed proximal part. The neuron expresses $Ca^{++}$ sensor - GCaMP6s.

**Movie S4.**

Instant sealing of an axon cut-end that does not let $Ca^{++}$ entry into DRG neuron through proximal part after axotomy. The neuron expresses $Ca^{++}$ sensor - GCaMP6s.

**Movie S5.**

Axotomy and shrinkage of a GFP-expressing DRG neuron imaged in 3D to visualize the volume change.

**Movie S6.**

Complete amputation of an axon of a DRG neuron during shrinkage after axotomy, which results in a very efficient sealing.
